# Autoinhibition of cMyBP-C by its middle domains

**DOI:** 10.1101/2024.07.19.603145

**Authors:** A.C. Greenman, R.L. Sadler, S.P. Harris

## Abstract

Cardiac myosin binding protein-C (cMyBP-C) is a sarcomere regulatory protein consisting of 11 well-folded immunoglobulin-like (Ig-like) and fibronectin type-III domains with the individual domains numbered C0-C10. Despite progress in understanding the functions of the N’ and C’-terminal ends of the protein, our understanding of the functional effects of the middle domains (C3-C4-C5-C6-C7) is still limited. Here we aimed to determine the functional significance of the middle domains by replacing endogenous cMyBP-C with recombinant proteins with and without the middle domains using our “cut and paste” SpyC3 mouse model. Specifically, we deleted domains C3-C7 or substituted these domains with unrelated Ig-like domains from titin to behave as inert “spacer” domains. Replacement with the spacer constructs resulted in a significant increase in myofilament calcium sensitivity, an almost instantaneous redevelopment of tension after a slack re-stretch protocol, and altered stretch activation responses, suggesting that the middle domains are functionally relevant and normally exert inhibitory effects on force development. We also investigated the significance of a flexible linker between domains C4 and C5 and a unique 28 amino acid loop insertion in C5. Whereas deletion of the C5 loop had no effect on force, deletion of the linker between C4 and C5 had comparable effects to deletion of domains C3-C7. Taken together, these data indicate that the middle domains play an important role in limiting the activating effects of the C0-C2 domains and that the C4C5 linker contributes to these effects.

**Significance Statement:** The functional role of the middle domains of cardiac myosin binding protein-C (cMyBP-C) are poorly understood, in part due to technical challenges inherent to *in vitro* methods that have mainly been used to study recombinant N’-terminal domains in the absence of the whole protein. Here we overcome this barrier by using a “cut and paste” approach, selectively removing and replacing the middle domains of cMyBP-C in permeabilized cardiomyocytes. Substituting the middle domains with titin Ig-like domains resulted in a large increase in myofilament calcium sensitivity, almost instantaneous redevelopment of force, and altered response to rapid stretch. Deletion of only the C4-C5 linker (11 amino acids) qualitatively resulted in the same alterations in force mechanics, albeit to a lesser magnitude. We suggest that the middle domains directly affect the regulation of cardiac muscle function by inhibiting the activating effects of the N’-terminal domains of cMyBP-C.

## Introduction

Cardiac myosin binding protein-C (cMyBP-C) is a key regulatory protein located in the C-zone of the sarcomere. cMyBP-C consists of 11 domains, eight immunoglobulin (Ig)-like and three fibronectin type-III III, a regulatory motif (i.e., the M-domain), and a proline/alanine-rich region between domains C0 and C1^1^. The N’-terminus (C0-C2) regulates muscle function by its interactions with both actin and myosin resulting in both activating and inhibitory effects on muscle function^1–3^. Mutations in *MYBPC3*, the gene encoding cMyBP-C, account for up to 50% of hypertrophic cardiomyopathy (HCM) cases with an identified genetic component^4^. Clinical presentations of HCM vary greatly, but is generally characterized by left ventricular hypercontractility, hypertrophy, myofibril disarray, and a higher risk of sudden cardiac death^5,6^. There are over 197 known HCM mutations occurring in *MYBPC3* alone^7^, and a majority of those mutants occur within the middle region of the protein, domains C3-C4-C5-C6-C7^8^.

However, much of the research effort to date has been spent on understanding the N’-terminal domains (NTD), generally considered C0, proline/alanine-rich region, C1, M-domain, and C2, of cMyBP-C (Figure 1A) in part because there is a growing consensus that the NTD dynamically regulate contraction and relaxation of the sarcomere. Specifically, the NTD bind to actin ^9–12^ and myosin ^13–18^ regulating muscle function by sensitizing the myofilaments to Ca^2+^ and/or by slowing cross-bridge kinetics^19–23^. Recent structural insights into the C’-terminal domains, generally considered C8, C9, and C10, have also been provided by cryo-electron microscopy (EM) and cryo-electron tomography (ET) reconstructions showing that these domains anchor the protein into its endogenous position on the thick filament^24–26^ and may bind to myosin heads.^27,28^

**Figure 1:**
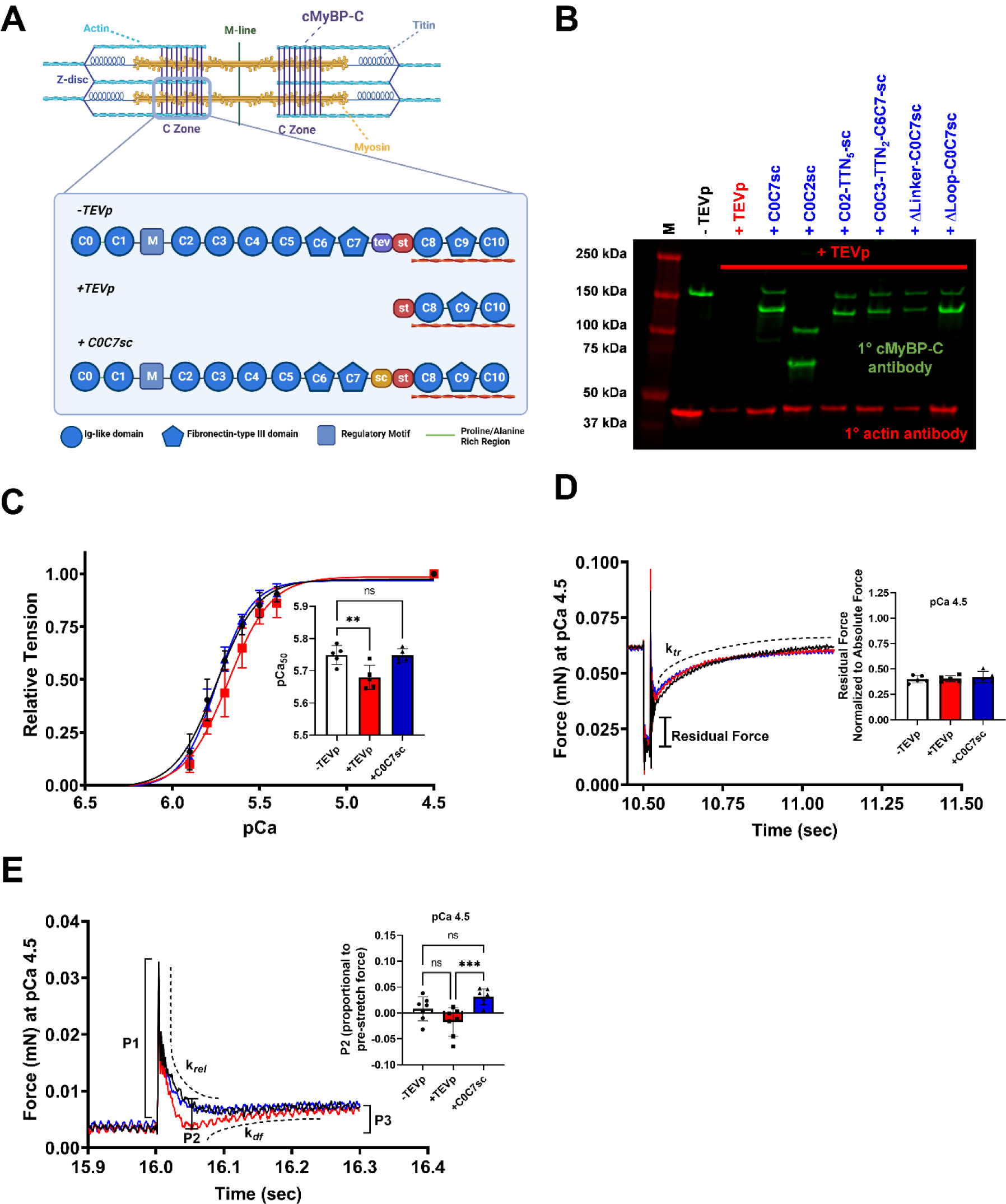
Effects of cut and paste of the C0-C7 domains of cMyBP-C on muscle mechanics. A) Cartoon representing the cut and paste SpyC3 system showing loss of endogenous C0-C7 domains after treatment with TEVp followed by replacement with recombinant C0-C7sc. Figure made using Biorender. B) Immunoblot results showing loss of C0-C7 domains after TEVp treatment (+ TEVp) and the replacement steps using several different recombinant proteins included in this study (e.g., + TEVp + C0C7sc). C) Normalized tension-pCa relationships before TEVp treatment (black), after treatment with TEVp (red), and after replacement with 20 µM C0C7sc (blue). D) Representative force traces of the slack re-stretch protocol for measurement of k*_tr_* and residual force. Inset shows summary data for residual force at pCa 4.5 for the three repeated conditions. E) Representative force trace of the rapid 2% stretch protocol for analysis of k*_rel_*, k*_df_*, P1, P2, and P3 amplitudes. Inset shows summary data for P2 amplitude across the three conditions, pCa 4.5. ** = p < 0.01, *** = p < 0.001

Comparatively less is known about the so-called middle domains of cMyBP-C, i.e., domains C3-C4-C5-C6-C7. Recent evidence suggests that these domains can bind to myosin S1 and S2 with high affinity, suggesting that the middle domains may help stabilize the parked conformation of myosin heads on the thick filament^16,29^.

There are also unique structures that occur within the middle domains. For example, the C4-C5 domains of cMyBP-C have a short ∼10 amino acid linker between C4 and C5 and a 28 amino acid insertion in C5, referred to as the C5 loop^30–32^. Using molecular dynamic simulations, Doh et al. showed that C4-C5 could form a bent and latched confirmation, which lead the authors to suggest that the C5 loop may help stabilize the bent confirmation of C4-C5^30^. Others have found that domains C6-C10 have high binding affinity for actin *in vitro*^33^ and domains C5-C10, but not C0-C4, restricted actin torsion^34^. However, the functional significance of middle domain binding interactions or a hinge-and-latch mechanism is unknown.

Until now, one of the challenges to studying the functional significance of the middle domains has been the prevalent use of shortened recombinant protein constructs *in vitro* that either lack the middle domains and/or the C’-terminal domains so that localization of the recombinant proteins cannot be restricted to the position of endogenous cMyBP-C in the C-zones. Here, we overcame this barrier by using a novel “cut and paste” approach to selectively remove and replace desired cMyBP-C domains in permeabilized myocytes *in situ*^20^. Using this method, we were able to remove (“cut”) endogenous C0-C7 domains and “paste” back a recombinant protein containing any desired modification of the middle domains. Using this approach, we show for the first time that the middle domains exert strong inhibitory effects that modulate effects of the N’-terminal domains to activate the thin filament.

## Methods

### SpyC3 Mice

Treatment of all animals was in strict accordance with the guidelines and protocols established by the Institutional Animal Care and Use Committee at the University of Arizona (protocol 13-446). SpyC3 mice (C57BL/6NJ background) were generated by CRISPR-Cas9 gene editing, as previously described^20^. Male and female SpyC3 mice were killed by cervical dislocation after anesthesia with isoflurane. Hearts were rapidly excised and dissected in ice-cold oxygenated Ringer’s buffer containing (in mM): 100 NaCl, 24 NaHCO_3_, 25 KCl, 2 MgSO_4_*7H_2_0, 1 Na_2_HPO_4_, and 0.2µM CaCl_2_. Left ventricles (including the interventricular septum) were frozen in liquid nitrogen and stored at –80°C until use.

### Cut and Paste Replacement of cMyBP-C

Homozygous SpyC3 mice have a tobacco etch virus protease (TEVp) consensus site and a SpyTag cassette inserted between domains C7 and C8 of cMyBP-C^20^. Removal of endogenous C0-C7 cMyBP-C domains (cMyBP-C^C0C7^) was accomplished by treatment of detergent-permeabilized cardiomyocytes with 20 µM TEVp at 22.5°C for 30 minutes and washout of the cleaved C0C7 fragment with a Relax buffer containing (in mM): 100 KCl, 10 imidazole, 2 EGTA, 5 MgCl_2_, 4 ATP (A2383, Sigma), and 1:100 HALT^TM^ protease inhibitor cocktail (78430, ThermoFisher) at pH 7.0. Recombinant cMyBP-C proteins encoding a SpyCatcher (sc) at their C’-termini were then covalently ligated to a SpyTag^35^ exposed at the N’-terminus of C8 by incubating detergent-permeabilized myocytes with 20 µM recombinant protein for 15 minutes at 15°C.

### Expression and Purification of Recombinant cMyBP-C Proteins

Purification of recombinant mouse cMyBP-C was accomplished using plasmids obtained from GenScript© and VectorBuilder© that were codon optimized for expression in bacteria. Small deletions (Loop and Linker) in mouse cMyBP-C wildtype sequences were accomplished using a Q5 Site-Directed Mutagenesis Kit (New England Biolabs, E0554). The following primers (Invitrogen) were used for to generate the ΔLinker and ΔLoop-C0C7sc constructs respectively: forward: CCGCCGAAAATTCATCTG, reverse: CATGAAGTGCAGTTTCGC, forward: AAGAAACTGCTGTGCGAAAC, and reverse: GGTCTTCTGCCACACAAC. Plasmids underwent bacterial transformation using *E. coli* and standard Luria Bertani (Sigma-Aldrich, MKCR7832) and Terrific Broth medium (Millipore, 71491) protocols for growth and auto-induction of transformed *E. coli*^20^. *E. coli* cells (BL21(DE3), New England Biolabs, C2527H) were spun down at 6,000 g after growth reached the stationary phase (OD 600 ∼3-6) and quick-frozen in liquid nitrogen. Pellet lysis was completed by homogenizing the pellet in B-PER^TM^ buffer (ThermoFisher, 78243), with protease inhibitors (2 µg/mL pepstatin, 20 µM PMSF, 2 µM E64), 0.01% 2-mercaptoethanol, adding lysozyme (40 mg/15 mL, Sigma, L-6876), and treated with DNase (50 units/mL) for further cell lysis and reducing viscosity of bacterial lysates, respectively. The supernatant protein fraction from cell lysis was isolated by incubating the supernatant in HisPur^TM^ Ni-NTA Resin (ThermoScientific, 88221) overnight, or for two hours, depending on the construct, and protein was eluted from the column. Some recombinant proteins were run on a size-exclusion column (LP BIOrad system; HiPrep^TM^ 26/60 Sephacryl® S-300 HR, Sigma-Aldrich) to improve purity (see supplemental materials for final proteins).

### Preparation of Membrane-Permeabilized Cardiomyocytes

Left ventricular samples were thawed in ice-cold Relax buffer containing 1% Triton-X 100 (VWR, CAS 9002-93-1), 0.1% saponin (Sigma Aldrich, 47036), and HALT^TM^ protease inhibitor cocktail. Small pieces of left ventricular myocardium (5-10 mg) were briefly homogenized using a Polytron PT 1200E device (Kinematica AG, Switzerland). The remaining slurry was rotated at 4°C for 25 minutes. Detergents were washed away by centrifuging at 2,500 g for 3 min and washed in relaxing buffer without detergents. Membrane-permeabilized cardiomyocytes were then visualized using an Olympus IX53 (Model IX-53, Olympus Instrument Co., Japan) inverted microscope and selected myocytes were glued to a force transducer and motor (Model 403A & 315C, Aurora Scientific Inc., Aurora, Ontario, Canada) using silicon glue (Marineland aquarium sealant). After curing for 30 minutes, the cardiomyocyte was lifted and moved into a relaxing solution of pCa 9.0, where pCa = (-log[Ca^2+^]).

### Force Measurements

Three consecutive tension-pCa curves were measured for each multicellular cardiomyocyte preparation: before TEVp treatment, after TEVp treatment, and after treatment with recombinant protein. Permeabilized myocytes were stretched to a sarcomere length of 2.3 µm and temperature was kept at 15°C for experimental protocols. Each tension-pCa curve consisted of slacking the cellular prep by 20%, or 0.8 original cell length (L_0_), after steady state force was obtained in each pCa buffer to quantify tension. The order of submaximal pCa’s (6.2 - 5.4) used for force protocols was randomized and tension generated at submaximal pCa’s was normalized to maximal Ca^2+^ activated force at pCa 4.5. The rate of force redevelopment (k*_tr_*) was measured by slacking myocytes and then stretching the cell to 1.05 of L_0_ and then to 1.00 L_0_ immediately after the slack test (2 ms). k*_tr_*, rate of force redevelopment, was quantified by fitting to a single exponential curve using MATLAB (R203a). Residual force was calculated as described previously^36^. Stretch activation responses were measured by subjecting the cardiomyocyte to a 2% stretch for 3 sec^37^. The rate of force decay (k*_rel_*) and the rate of delayed force redevelopment (k*_df_*) were fitted to a single exponential curve. Each phase of the stretch activation protocol was normalized to the pre-stretch force.

### Immunoblotting and antibodies

Samples were prepared for immunoblotting by homogenizing left ventricular (LV) tissue (∼50 mg) in ice-cold Relax buffer containing 1% Triton, 0.1% Saponin, and HALT^TM^ protease inhibitor cocktail. The remaining wash, TEVp treatment (12 µg per mg tissue), and recombinant protein steps were done as previously described with permeabilized cardiomyocyte preps for force measurements^20^. The final homogenates were mixed with an equal volume of urea buffer ([in mol/L]: 8 urea, 2 thiourea, 0.05 Tris-HCl, 0.075 dithiothreitol with 3% SDS and 0.03% bromophenol blue, pH 6.8). Protein lysate samples were run on an SDS -PAGE gradient gel (4561086, 4-15% Mini-PROTEAN TGX^TM^ Precast Protein Gel, Bio-Rad) and transferred onto a nitrocellulose membrane, as previously described^38^. Blots were blocked with OneBlock^TM^ Fluorescent Blocking Buffer (20-314, Genessee Scientific) and probed with antibodies to custom primary rabbit polyclonal antibody to cMyBP-C (diluted 1:75,000, previously described^21^) and to a mouse monoclonal antibody to F-actin (diluted 1:2,000, Invitrogen, clone ACTN05 C4) as a loading control. Secondary antibodies were diluted 1:10,000 (IRDye 800CW goat anti-rabbit and 680RD goat anti-mouse). Blots were imaged on an Odyssey CLx system.

### Statistical Analyses

All values are reported as mean ± SD. Repeated measures one-way ANOVA with multiple comparisons was used to compare control measurements to measurements after TEVp treatment and following recombinant protein replacements. If values were missing, a mixed effects test was used to determine differences between treatment groups. When only two of the three conditions were compared (see no fit (NF) data), a paired t-test was used. Significance was considered at p < 0.05.

## Results

### Cut and paste of domains C0-C7 reversibly alters myocyte force and crossbridge kinetics

Figure 1A-B depicts a diagram and representative western blot images using the cut and paste method. Treatment of permeabilized cardiomyocytes with TEVp cleaved endogenous C0C7 (cMyBP-C^C0C7^) which was removed from the sarcomere by washing. Next, recombinant proteins were added back to permeabilized cell lysates and ligated (“pasted”) proteins were visualized using an antibody to cMyBP-C^39^ (Figure 1B).

Removal of domains C0-C7 (cMyBP-C^C0C7^) following TEVp treatment led to a significant rightward shift in the tension-pCa relationship compared to before TEVp treatment (pCa_50_ 5.75 ± 0.03 vs. 5.68 ± 0.04, p = 0.001, Figure 1C and Table I) and faster rates of force re-development and decay (k*_tr_*) as previously reported^20^. Loss of cMyBP-C^C0C7^ also altered responses to stretch by significantly accelerating k*_rel_*. The P2 amplitude tended to be reduced following TEVp treatment but was not significantly different compared to before TEVp levels (p = 0.09). As expected, ligation with recombinant C0C7sc reversed all effects of cMyBP-C^C0C7^ loss, causing a leftward shift of the tension-pCa relationship and slowing k*_tr_* and k*_rel_* back to control values prior to TEVp treatment (Fig. 1 and Table I). P2 amplitude was also significantly greater with pasting of wildtype recombinant C0C7sc compared to post-TEVp treatment, but not compared to baseline conditions prior to TEVp treatment (p = 0.0005 and p = 0.077, respectively). However, we noted there was considerable variability in achieving good fits of an exponential equation to k*_df_* curves at pCa 4.5 (maximal Ca^2+^). Despite this limitation, we found that k*_df_* was not different between baseline and after pasting C0C7sc at an intermediate Ca^2+^ activation near the pCa_50_ of force activation (pCa 5.7, supplementary materials Fig. S4).

**Table I.** Effects of C0C7sc, C5C7sc, C3C7sc, and C0C2sc on force and crossbridge kinetics.

| Conditions |  | Active Tension<br>(mN/mm<sup>2</sup>) | Passive Tension<br>(mN/mm<sup>2</sup>) | K<sub>tr</sub> at pCa 4.5<br>(sec<sup>-1</sup>) | K<sub>rel</sub> at pCa 4.5<br>(sec<sup>-1</sup>) | P3 amplitude<br>(normalized to<br>pre-stretch<br>force, pCa 4.5) |
| --- | --- | --- | --- | --- | --- | --- |
| C0C7sc | Before TEVp | 15.92 ± 4.86 | 2.00 ± 0.86 | 8.97 ± 0.97 | 141.13 ± 32.8 | 0.038 ± 0.01 |
|  | After TEVp | 17.12 ± 6.09 | 1.53 ± 0.89 | 10.61 ± 1.30‡ | 184.67 ± 46.4‡ | 0.038 ± 0.008 |
|  | C0C7sc | 19.12 ± 6.27 | 2.32 ± 0.76 | 9.02 ± 1.33 | 159.93 ± 58.7 | 0.042 ± 0.006 |
| C5C7sc | Before TEVp | 20.27 ± 10.82 | 2.39 ± 1.46 | 8.56 ± 1.64 | 60.04 ± 19.3 | 0.031 ± 0.005 |
|  | After TEVp | 22.33 ± 10.76 | 2.13 ± 1.52 | 9.79 ± 1.40‡ | 82.44 ± 22.87‡‡ | 0.032 ± 0.007 |
|  | C5C7sc | 22.69 ± 11.74 | 1.99 ± 1.39 | 9.16 ± 1.26 | 68.57 ± 25.0 | 0.036 ± 0.009 |
| C3C7sc | Before TEVp | 17.63 ± 10.18 | 2.10 ± 1.24 | 12.30 ± 1.07 | 78.48 ± 5.36 | 0.035 ± 0.012 |
|  | After TEVp | 15.77 ± 8.40 | 0.96 ± 0.61 | 14.31 ± 1.28‡ | 97.23 ± 10.77‡‡ | 0.035 ± 0.009 |
|  | C3C7sc | 14.96 ± 9.47 | 0.97 ± 0.37 | 12.26 ± 1.93 | 83.24 ± 5.42 | 0.036 ± 0.015 |
| C0C2sc | Before TEVp | 15.99 ± 6.11 | 1.47 ± 0.55 | 9.29 ± 0.83 | 142.60 ± 32.28 | 0.034 ± 0.010 |
|  | After TEVp | 15.74 ± 6.48 | 1.18 ± 0.39‡ | 11.14 ± 1.36‡‡ | 176.02 ± 63.16 | 0.034 ± 0.008 |
|  | C0C2sc | 16.09 ± 6.04 | 2.29 ± 0.81 | NF | 119.75 ± 9.49 | 0.060 ± 0.018* |

### The N’-terminal domains C0-C2 alone only partially restored functional effects after loss of cMyBP-C^C0C7^

To determine whether the N’-terminal domains alone are sufficient to recover force and kinetics after loss of cMyBP-C^C0C7^, we next pasted back a shorter recombinant protein containing only the C0-C2 domains of cMyBP-C lacking the middle domains C3-C7. Similar to full-length C0C7sc, ligation of C0C2sc caused a leftward shift of the tension-pCa relationship that rescued the rightward shift following TEVp treatment and resulted in a small but significant increase in pCa_50_ compared to the control curve prior to TEVp treatment (Fig. 2; pCa_50_ 5.71 ± 0.04 vs. 5.76 ± 0.04). However, C0C2sc also significantly altered the k*_tr_* and stretch activation responses in a manner different than C0C7sc. As shown in Fig. 2B, there was an almost instantaneous redevelopment of steady state tension after a slack re-stretch protocol resulting in a high level of “residual” force that prevented accurate curve fits to obtain k*_tr_* values. After C0C2sc replacement, the residual force, or the difference in force between the end of the re-stretch maneuver and the slack force, was significantly greater compared to either before or after TEVp treatment (p = 0.0008 and p = 0.0003, respectively, Fig. 2B inset). The response to a rapid 2% stretch was also altered by C0C2sc, as shown in Fig. 2C. Specifically, the amplitude of P2, normalized to pre-stretch force, was significantly greater after ligation with C0C2sc compared to before TEVp treatment (p = 0.02, Fig. 2C inset). The rate of force decay following the initial stretch was not different compared to the before TEVp condition (p = 0.3). The amplitude of P3 was increased (p = 0.02 compared to before TEVp, Table I), presumably due to the increased P2 amplitude. As with k*_tr_* measurements, there was no observable k*_df_* segment of the response, making it difficult to accurately fit an exponential curve after pasting C0C2sc (Fig. 2C). Taken together, these data indicate that domains C0-C2, like C0C7sc, have Ca^2+^ sensitizing effects on steady state force, but that C0C2sc and C0C7sc have different effects on crossbridge kinetics.

**Figure 2:**
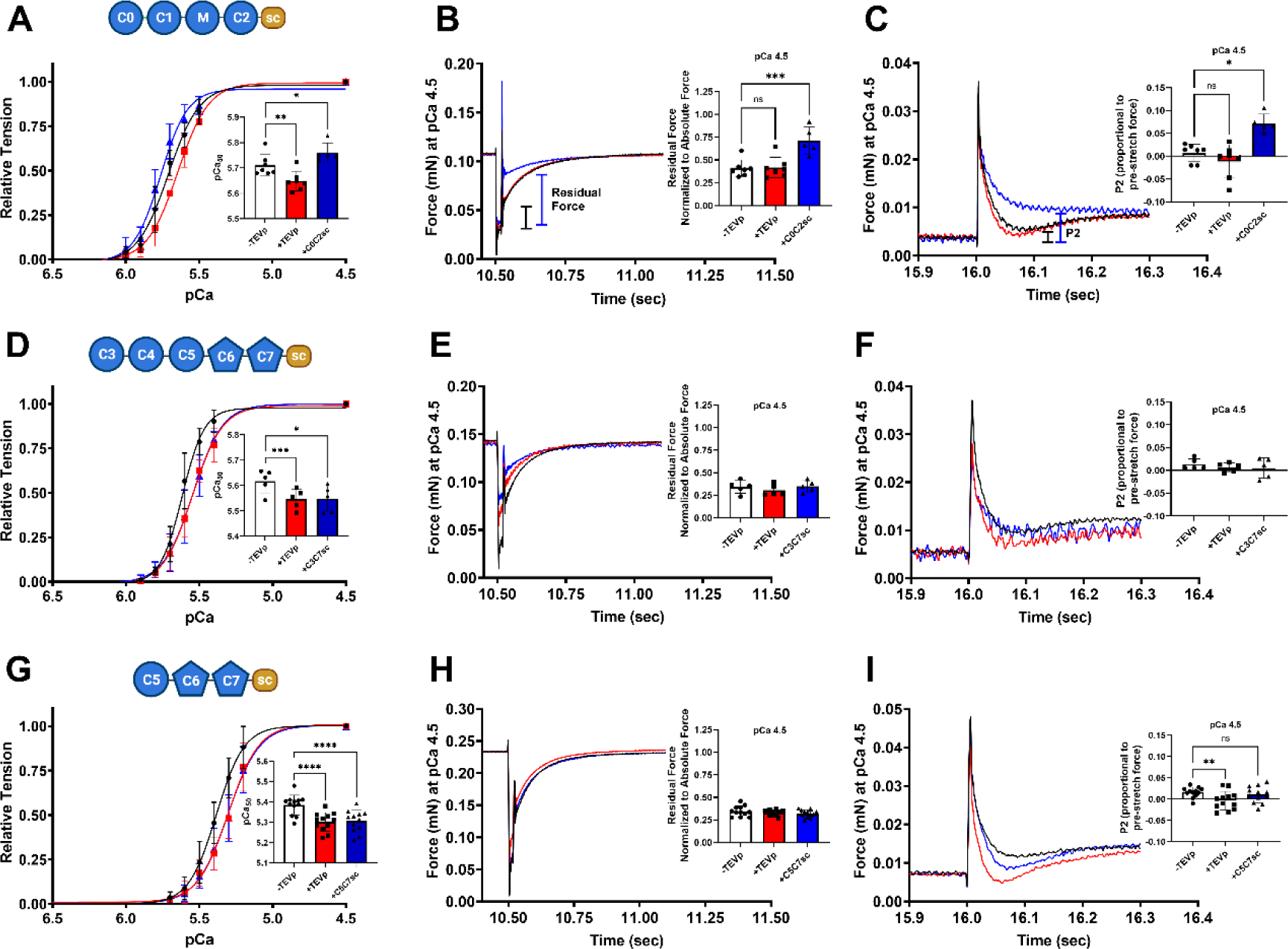
Functional Effects of replacing cMyBP-C^C0C7^ with C5C7sc, C3C7sc, and C0C2sc. A-C) Normalized tension-pCa relationships, representative force traces of the slack re-stretch protocol, and rapid 2% stretch protocol for before TEVp (black), after 20 µM TEVp (red), and after replacement with 20 µM C5C7sc (blue). D-F) Normalized tension-pCa relationship, representative force traces of the slack re-stretch protocol, and rapid 2% stretch protocol for before TEVp (black), after 20 µM TEVp (red), and after replacement with 20 µM C3C7sc (blue). G-I) Normalized tension-pCa relationships, representative force traces of the slack re-stretch protocol, and rapid 2% stretch protocol for before TEVp (black), after 20 µM TEVp (red), and after replacement with 20 µM C0C2sc (blue). * = p < 0.05, ** = p < 0.01, *** = p < 0.001, **** = p < 0.0001.

### C3C7sc and C5C7sc alone do not affect force or crossbridge kinetics

Based on the differences observed between C0C2sc and C0C7sc, we next sought to determine whether the middle domains contribute directly to the functional effects of cMyBP-C by replacing cMyBP-C^C0C7^ with recombinant proteins containing only the middle domains, i.e., C3C7sc or C5C7sc (recombinant proteins lacking NTD, C0C2). However, as shown in Fig. 2, neither C3C7sc nor C5C7sc had any discernable effects on myofilament Ca^2+^ sensitivity of tension (Figure 2D&G). Notably, cross-bridge kinetics sped up with TEVp treatment (k*_tr_* and k*_rel_*, Table I), but pasting C3C7sc and C5C7sc led to a recovery of k*_tr_* and k*_rel_* comparable to pre-TEVp treatment levels. The amplitude of P2 of stretch activation was significantly reduced with TEVp treatment (p = 0.005, Figure 2I) and recovered to baseline levels when pasting back C5C7sc. Replacement with C3C7sc did not significantly alter the P2 amplitude when compared to before and after TEVp conditions (Figure 2F). k*_df_* was not different between baseline and after pasting C3C7sc or C5C7sc at submaximal calcium (supplementary materials Fig. S4) Taken together, these results suggest that the middle domains alone do not directly affect steady state tension or crossbridge kinetics, but that when they are present in C0C7sc they have modulatory effects on C0C2.

### Replacing domains C3-C7 with titin Ig-like domains exaggerates effects of C0C2sc

To determine whether the effects of the C3-C7 domains in combination with C0-C2 are specific to cMyBP-C or whether these 5 middle domains could be replaced with any similarly folded Ig-like domains comparable in size, we next replaced domains C3-C7 with 5 unrelated Ig-like domains (Ig27 from titin) in the recombinant protein C0C2-TTN_5_-sc. Importantly, because deletion of C3-C7 (as in C0C2sc) is expected to shorten the overall contour length of cMyBP-C by the length of 5 Ig domains, the length of C0C2-TTN_5_-sc should be approximately similar in length to C0C7sc. C0C2-TTN_5_-sc should therefore help distinguish whether the functional differences observed between C0C2sc and C0C7sc are due to specific effects of the C3-C7 domains or instead to other non-specific factors such as protein length. As shown in Fig. 3, ligation of C0C2-TTN_5_-sc unexpectedly resulted in a dramatic increase in Ca^2+^ sensitivity of tension relative to baseline (control) values before TEVp treatment (ΔpCa_50_ = 0.26 ± 0.04, p = 0.003, Fig. 3A). Notably, this shift in pCa_50_ far exceeded that induced by C0C2sc alone (Fig. 2). Tension at low submaximal concentrations of Ca^2+^ (e.g., pCa 6.2 – 5.7) were disproportionately increased causing a large reduction in the Hill slope (p = 0.004, Supplementary Table SI). However, similar to C0C2sc, ligation of C0C2-TTN_5_-sc led to an almost instantaneous redevelopment of tension after a slack-re-stretch protocol, resulting in a large increase in residual force compared to before TEVp (p = 0.0005, Fig. 3B). C0C2-TTN_5_-sc also caused an exaggerated stretch activation response when compared to before and after TEVp treatment (Fig. 3C & Table II). Ligation of C0C2-TTN_5_-sc led to a reduction in k*_rel_* compared to TEVp treatment (Table II). Similar to effects of C0C2sc, k*_df_* curves after pasting C0C2-TTN_5_-sc were minimal and difficult to fit accurately (Fig. 3C). These results show that effects of C0C2-TTN_5_-sc are qualitatively similar to those of C0C2sc, but are much greater in magnitude.

**Figure 3:**
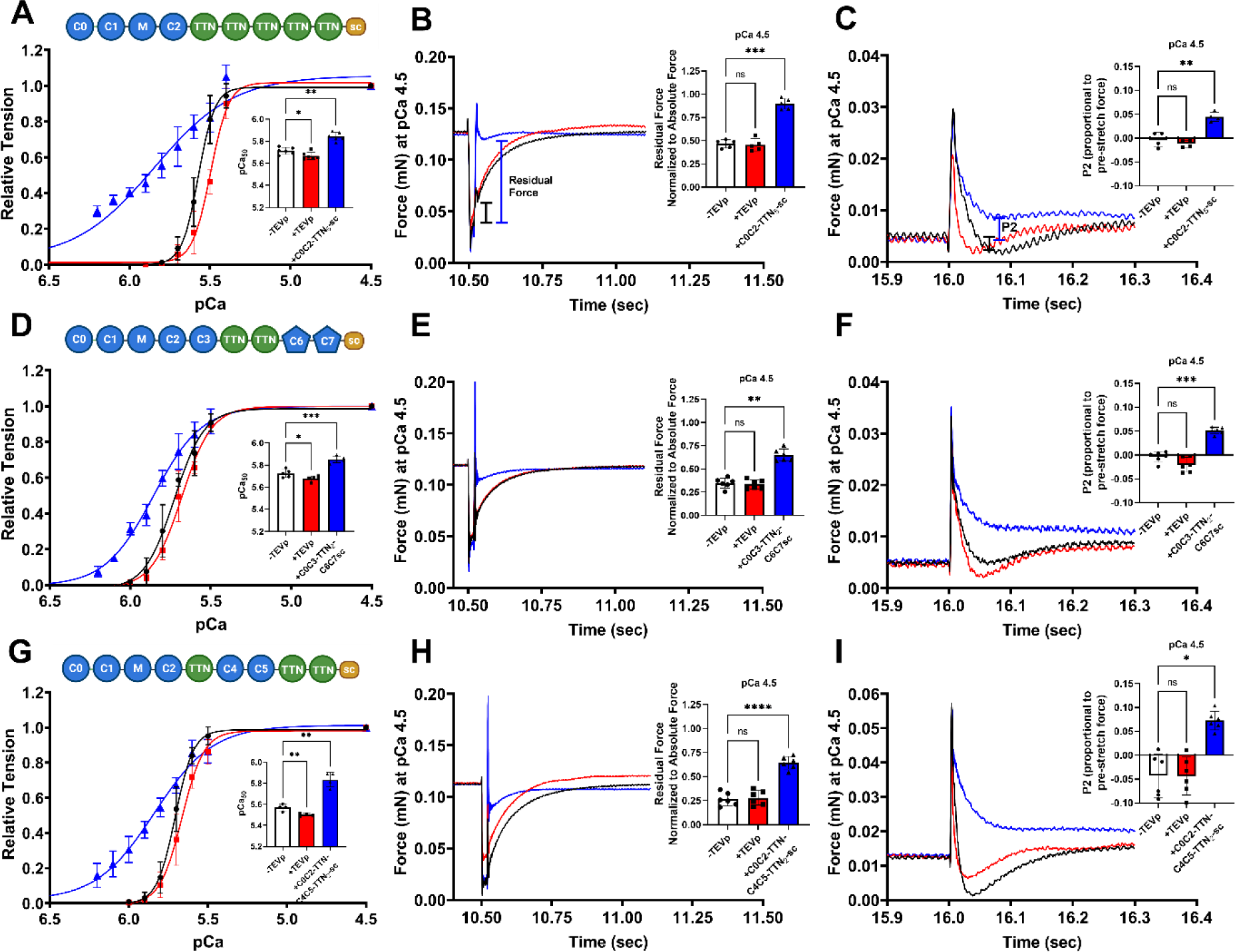
Functional effects of recombinant TTN-sc proteins. A-C) Normalized tension-pCa relationship, representative force traces of the slack re-stretch protocol, and rapid 2% stretch protocol for before TEVp (black), after 20 µM TEVp (red), and after replacement with 20 µM C0C2-TTN_5_-sc (blue). D-F) Normalized tension-pCa relationship, representative force traces of the slack re-stretch protocol, and rapid 2% stretch protocol for before TEVp (black), after 20 µM TEVp (red), and after replacement with 20 µM C0C3-TTN_2_-C6C7-sc (blue). G-I) Normalized tension-pCa relationship, representative force traces of the slack re-stretch protocol, and rapid 2% stretch protocol for before TEVp (black), after 20 µM TEVp (red), and after replacement with 20 µM C0C2-TTN-C4C5-TTN_2_-sc (blue). * = p < 0.05, ** = p < 0.01, *** = p < 0.001, **** = p < 0.0001.

**Table II.**
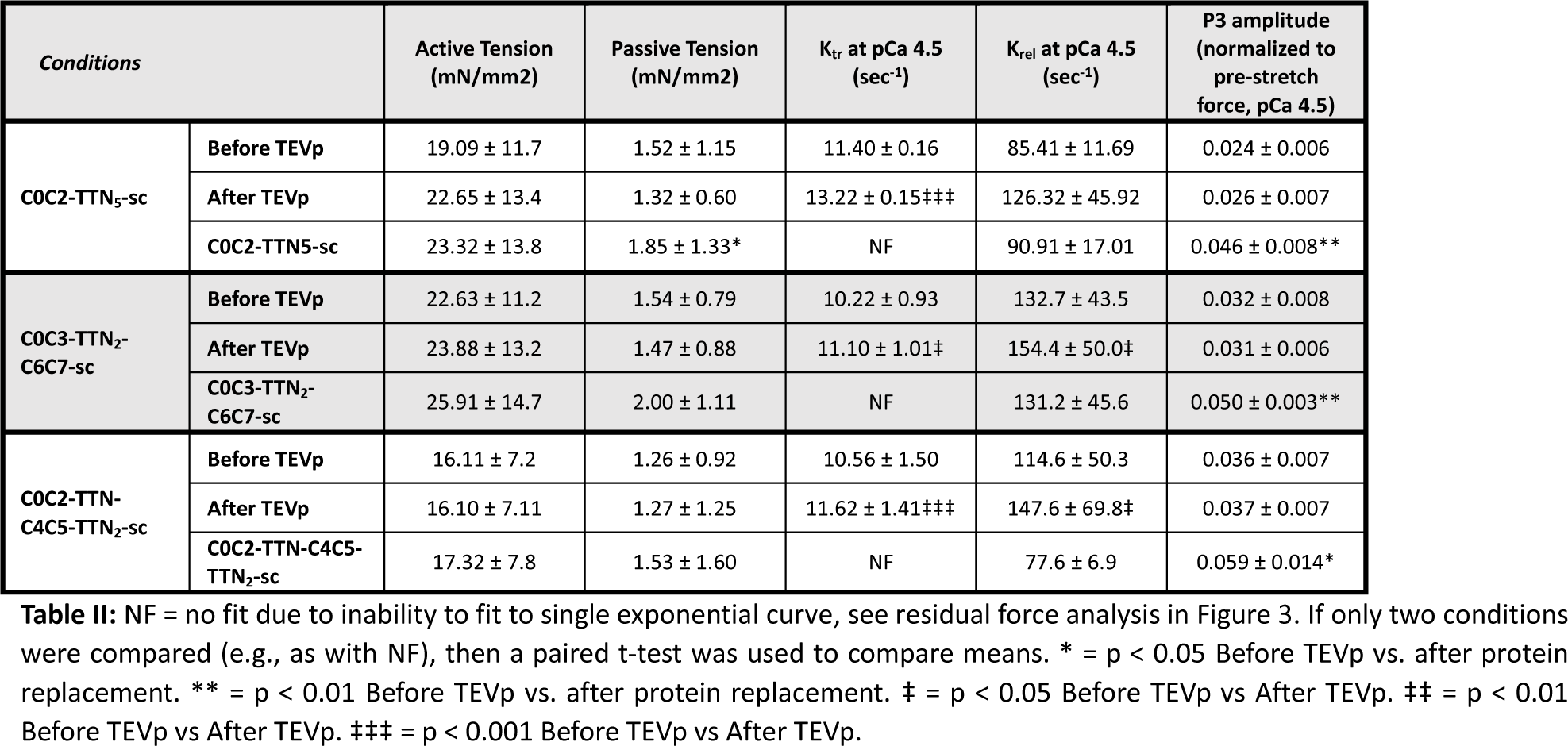
Effects of recombinant TTN-sc proteins on force and crossbridge kinetics.

To test the possibility that the substituted TTN domains directly affected force and crossbridge kinetics, we also replaced cMyBP-C^C0C7^ with recombinant TTN_5_-sc (without any cMyBP-C domains). Results shown in Supplemental Fig. 2 demonstrate that the 5 TTN domains alone had no effect on Ca^2+^ sensitivity of tension or on crossbridge cycling kinetics as assessed by k*_tr_* and transient stretch activation responses, as expected if these domains do not exert any functional effects on their own. Taken together, we conclude that C3-C7 domains of cMyBP-C exert specific modulatory effects on cMyBP-C NTD (C0-C2) that are not duplicated by unrelated Ig domains of similar size.

### Selective domain substitutions reveal that all 5 middle domains likely contribute to the modulatory effects of C3-C7

To narrow down which of the 5 middle domains are necessary to modulate the functional effects of C0C2, we next replaced only selected cMyBP-C middle domains with TTN Ig domains. Because domains C4 and C5 have unique structural features that were recently suggested to work together in concert (i.e., the flexible linker between C4-C5 and the unstructured loop in C5),^30^ we first deleted C4 and C5 and replaced only these cMyBP-C domains with TTN Ig domains. Functional effects of the resulting recombinant protein, C0C3-TTN_2_-C6C7-sc, are shown in Fig. 3. This C4-C5 substitution mutant had qualitatively similar effects to the 5-domain substitution protein, C0C2-TTN5-sc, but the magnitude of effects was reduced compared to C0C2-TTN_5_-sc. For example, the leftward shift in pCa_50_ produced by C0C3-TTN_2_-C6C7-sc compared to before TEVp treatment was still significant but reduced in magnitude (ΔpCa_50_ = 0.12 ± 0.03, p = 0.002, Fig. 3D) compared to C0C2-TTN_5_-sc. Similarly, residual force, P2, and P3 amplitudes were also significantly increased with C0C3-TTN_2_-C6C7-sc, albeit to a lesser extent than C0C2-TTN_5_-sc (Fig. 3E-F & Table II). As for C0C3-TTN_2_-C6C7-sc, the k*_df_* curve was difficult to fit after pasting C0C3-TTN_2_-C6C7-sc (Fig. 3F). These data indicate that domains C4 and C5 are necessary to modulate C0-C2 function but alone they are not sufficient to account for all of the regulatory effects of the middle domains of cMyBP-C.

Because the C4-C5 substitution protein (C0C3-TTN_2_-C6C7-sc) did not fully account for functional differences between C0C7sc and C0C2-TTN_5_-sc, we next substituted the remaining cMyBP-C domains, C3, C6, and C7 with TTN domains while leaving C4 and C5 in place. Functional effects of the resulting recombinant protein, C0C2-TTN-C4C5-TTN_2_-sc, are shown in Fig. 3. Similar to effects of deletion and replacement of C4-C5 with unrelated titin Ig-like domains, substitution of C3, C6, and C7 resulted in intermediate effects between C0C7sc and C0C2-TTN_5_-sc. Taken together, these data indicate that all 5 middle domains of cMyBP-C (C3-C7) likely contribute to modulation of effects of the N’-terminal domains C0-C2.

### Deletion of the C4-C5 Linker, but not the C5 Loop, mimics substitution of the C4-C5 domains

Finally, as an additional test of the hypothesis that the C4-C5 linker and the C5 unstructured loop regulate cMyBP-C function via a hinge-and-latch mechanism^30^, we next deleted either the C4-C5 linker (ΔLinker-C0C7sc, deletion of 11 amino acids between C4 and C5) or the C5 loop (ΔLoop-C0C7sc, deletion of 28 amino acids within C5)^30^. As shown in Fig. 4, deletion of the C4-C5 linker increased pCa_50_, residual force, P2, and P3 amplitudes compared to baseline values (before TEVp treatment) (Fig, 4A-C & Table III). Notably, the ΔpCa_50_ induced by the ΔLinker-C0C7sc and C0C3-TTN_2_-C6C7-sc recombinant proteins were not significantly different (ΔpCa_50_ = 0.10 ± 0.03 & 0.12 ± 0.03, respectively, p = 0.3). However, ligation of ΔLoop-C0C7sc led to a complete rescue of pCa_50_ to before TEVp levels (ΔpCa_50_ = -0.001 ± 0.01, Fig. 4D) and there were no other differences in residual force, P2, or P3 between ΔLoop-C0C7sc and before TEVp treatment (Fig. 4E-F & Table III). Reduced residual force allowed k*_tr_* values to be calculated after pasting ΔLoop-C0C7sc, in contrast to ΔLinker-C0C7sc results (Table III), and k*_tr_* was not different before TEVp compared to pasting ΔLoop-C0C7sc. k*_df_* curve fits were also possible after pasting with ΔLoop-C0C7sc (Fig4C&F and supplementary Fig. S4), but not after pasting with ΔLinker-C0C7sc. These data indicate that the C4-C5 linker, but not the C5 loop, is necessary for the modulatory effects of C3-C7 on the N’-terminal domains of cMyBP-C.

**Figure 4:**
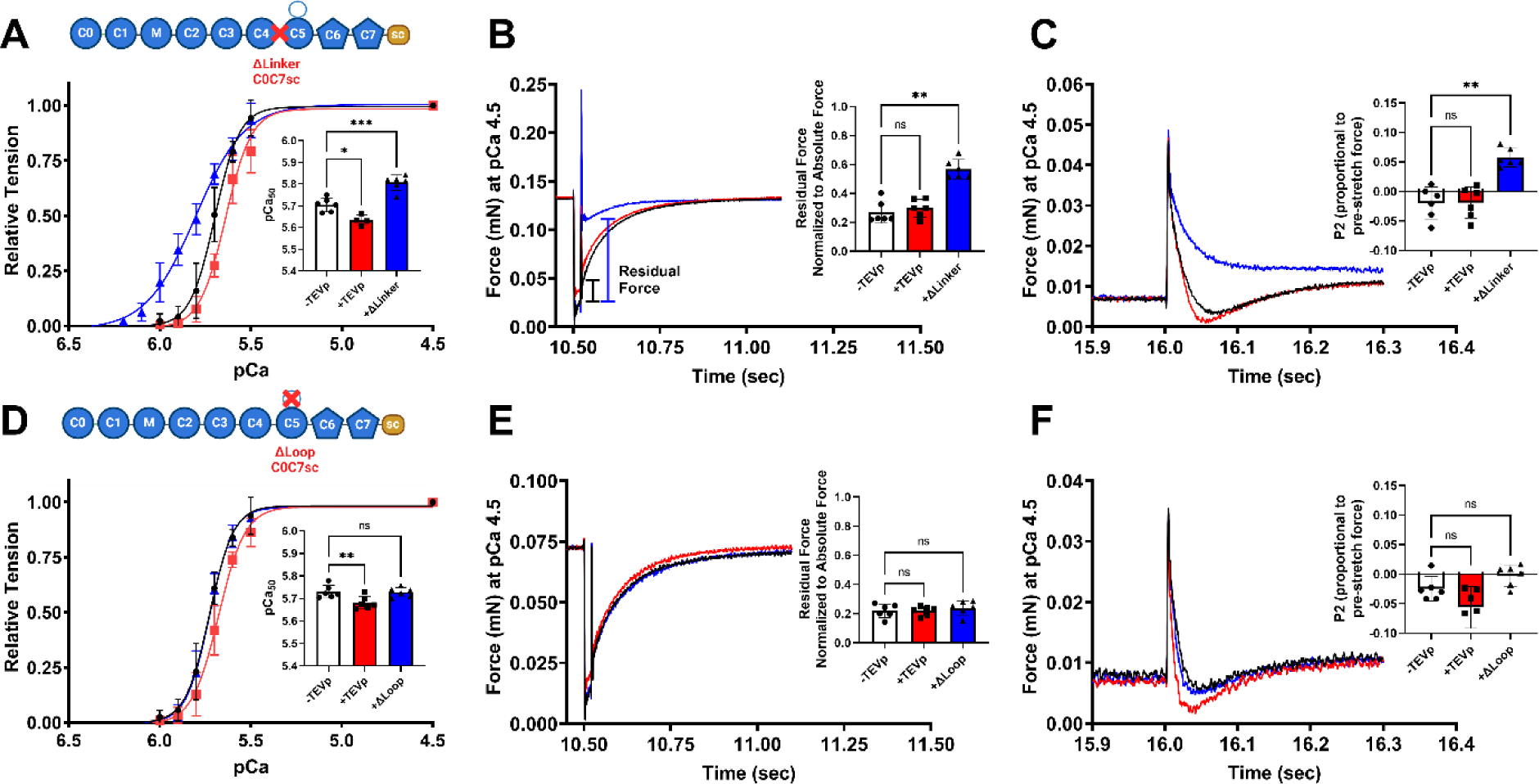
Functional effects of deleting either the C4C5 Linker or the C5 Loop. A-C) Normalized tension-pCa relationship, representative force traces of the slack re-stretch protocol, and rapid 2% stretch protocol for before TEVp (black), after 20 µM TEVp (red), and after replacement with 20 µM ΔLinker C0C7sc (blue). D-F) Normalized tension-pCa relationship, representative force traces of the slack re-stretch protocol, and rapid 2% stretch protocol for before TEVp (black), after 20 µM TEVp (red), and after replacement with 20 µM ΔLoop C0C7sc (blue). * = p < 0.05, ** = p < 0.01, *** = p < 0.001.

**Table III.** Effects of deleting the linker and loop on force and crossbridge kinetics.

| | ΔLinker C0C7sc | | | ΔLoop C0C7sc | | |
| --- | --- | --- | --- | --- | --- | --- |
| | Before TEVp | After TEVp | ΔLinker C0C7sc | Before TEVp | After TEVp | ΔLoop C0C7sc |
| Active Tension (mN/mm<sup>2</sup>) | 22.1 ± 7.9 | 23.3 ± 11.0 | 32.8 ± 16.1 | 20.6 ± 11.7 | 20.7 ± 9.9 | 22.1 ± 11.0 |
| Passive Tension (mN/mm<sup>2</sup>) | 1.73 ± 1.3 | 1.73 ± 1.0 | 2.5 ± 2.1 | 1.8 ± 1.1 | 1.9 ± 1.1 | 2.1 ± 1.3 |
| K<sub>tr</sub> | 11.9 ± 1.7 | 13.3 ± 2.3‡ | NF | 12.02 ± 2.94 | 13.32 ± 3.08 | 11.95 ± 2.40 |
| K<sub>rel</sub> | 119.4 ± 46.3 | 146.4 ± 44.7 | 96.7 ± 45.8 | 125.7 ± 17.4 | 171.7 ± 30.2‡ | 128.7 ± 25.1 |
| P3 | 0.034 ± 0.007 | 0.033 ± 0.007 | 0.061 ± 0.013** | 0.04 ± 0.008 | 0.05 ± 0.017 | 0.05 ± 0.006 |

## Discussion

The major finding from this study is that the middle domains of cMyBP-C (i.e., domains C3-C4-C5-C6-C7) exert specific functional effects within the context of full-length cMyBP-C. This was most evident when domains C3-C7 were replaced with 5 unrelated titin Ig domains in the recombinant protein C0C2-TTN_5_-sc. The replacement resulted in significant functional differences compared to full-length wildtype C0C7sc (with domains C3-C7 intact) including a profound increase in Ca^2+^ sensitivity of tension, high residual force after a release and re-stretch maneuver (k*_tr_*) and altered stretch activation responses. The significant augmentation of submaximal force in the absence of C3-C7 suggests that these domains normally exert inhibitory effects when present in the full-length protein. Results from the titin substitution experiments were qualitatively similar to effects in C0C2sc (lacking domains C3-C7), although effects on Ca^2+^ sensitivity of tension were greater in C0C2-TTN_5_-sc. It is unclear why there should be a difference in the magnitude of the shift in Ca^2+^ sensitivity of tension, but steric constraints due to the shorter contour length of C0C2sc compared to C0C2-TTN_5_-sc may play a role. Consistent with this, when C0C2 is added free in solution without SpyCatcher (which is necessary to anchor it the thick filament in SpyC3 myocytes), C0C2 causes a more profound leftward shift in Ca^2+^ sensitivity and a reduction in Hill slope similar to effects observed here with C0C2-TTN_5_-sc^9^.

The inhibitory effects of C3-C7 appear dependent on the presence of the N’-terminal domains, C0-C2. This is because recombinant C3C7sc (lacking C0-C2 domains) on its own had no discernable effects on force, arguing against direct effects of these domains on actin-myosin interactions. Yet, deletion or substitution of these domains in recombinant proteins containing C0-C2 had significant effects. One possibility is that when expressed in the full-length native protein domains C3-C7 play an autoinhibitory role that limit the effects of C0-C2 and/or promotes unbinding of C0-C2 from its sarcomere targets. For instance, C0-C2 domains have long been known to bind to both thick and thin filament targets where binding to actin and tropomyosin on the thin filament cause tropomyosin to shift towards its on state^39,40^, while C0-C2 promotes the parked interacting heads motif (IHM) conformation on the thick filament presumably through interactions with myosin S1, S2, and RLC ^16,41^. Recently, C3-C7 were also shown to bind to myosin S1 and S2 with high affinity leading to the suggestion that the middle domains may also directly stabilize myosin heads on the thick filament^29^. However, other binding partners for the middle domains were reported, including actin^33^ and inter-domain interactions, such as between C5 and C8^42^.

Our C3C7sc results are in broad agreement with other studies that demonstrated no change in pCa_50_ and k*_df_* after injecting knockout cMyBP-C mice with AAV9 vectors containing either N’-terminal truncated cMyBP-C (i.e., domains C3-C10 only) or full-length cMyBP-C^43^. However, there was full recovery of k*_rel_* and partial recovery of LV mass and global longitudinal strain with the C3C10 AAV9 group, leading the authors to suggest that both the N’-terminal domains and the middle domains of cMyBP-C are important for *in vivo* cardiac function^43^. We also found a small functional effect of C3-C10 on their own in permeabilized myocytes (k*_tr_* and k*_rel_*), supporting the functional role of C3-C7 in the intact, native protein.

Deletion of C3-C7 (as in C0C2-TTN_5_-sc) could in principle augment force and crossbridge cycling through either destabilization of myosin heads or enhanced activation of the thin filament. Interestingly, our data suggest that all 5 middle domains contribute to these effects because substitution of either C4 and C5, or C3, C6, and C7 with titin domains (C0C3-TTN_2_-C6C7-sc and C0C2-TTN-C4C5-TTN_2_-sc) resulted in intermediate effects compared to substitution of all 5 domains in C0C2-TTN_5_-sc. A graded effect was somewhat unexpected in part due to the focus on unique structural features present in C4 and C5, namely a flexible linker between domains C4 and C5 and a large 28 amino acid loop in C5^30^. Recent molecular dynamic simulations found that C4-C5 domains were able to form a tight V-like configuration only in constructs containing the 10-11 amino acid linker sequence, leading the authors to propose a hinge-and-latch mechanism where the V-shape formed by the C4-C5 linker is stabilized by the C5 loop^30^. Our data support a functional role for the C4-C5 linker because deletion of this short 11 amino acid sequence (ΔLinker-C0C7sc) caused comparable functional effects to substitution of domains C4 and C5 with titin domains. By contrast, deletion of the C5 loop (ΔLoop-C0C7sc) had no discernable effects on force because effects of ΔLoop-C0C7sc were not different from effects of wildtype C0C7sc in our assays.

The idea that the C4-C5 linker functions as a hinge^30,44,45^ suggests a plausible hypothesis for our results that the functional effects of the middle domains are dependent on the presence of C0-C2. For instance, if the C4-C5 hinge affects steric positioning of N’-terminal domains near their binding targets, then modulation of the linker could promote or inhibit these interactions. A scenario for C0-C2 binding to actin is shown in Fig. 5. In the case of C0-C2 binding to the thin filament, results with wildtype C0C7sc compared to ΔLinker-C0C7sc indicate that the C4-C5 linker may play a critical role in *unbinding* NTD from actin, i.e., moving C0-C2 away from the thin filament and thereby limiting thin filament activation. If so, mechanical effects of conformational changes in the linker could be critical in modulating diastolic relaxation. Similarly, a hinge mechanism may also explain the graded dependence of the functional effects, for example if all 5 middle domains interact with the N’-terminal domains to stabilize the bent *versus* extended hinge conformation. While there is not yet direct evidence for middle domain interactions with C0-C2, there is evidence for interdomain interactions within cMyBP-C (e.g., C5 and C8)^42^ and β-MyHC^46^.

**Figure 5:**
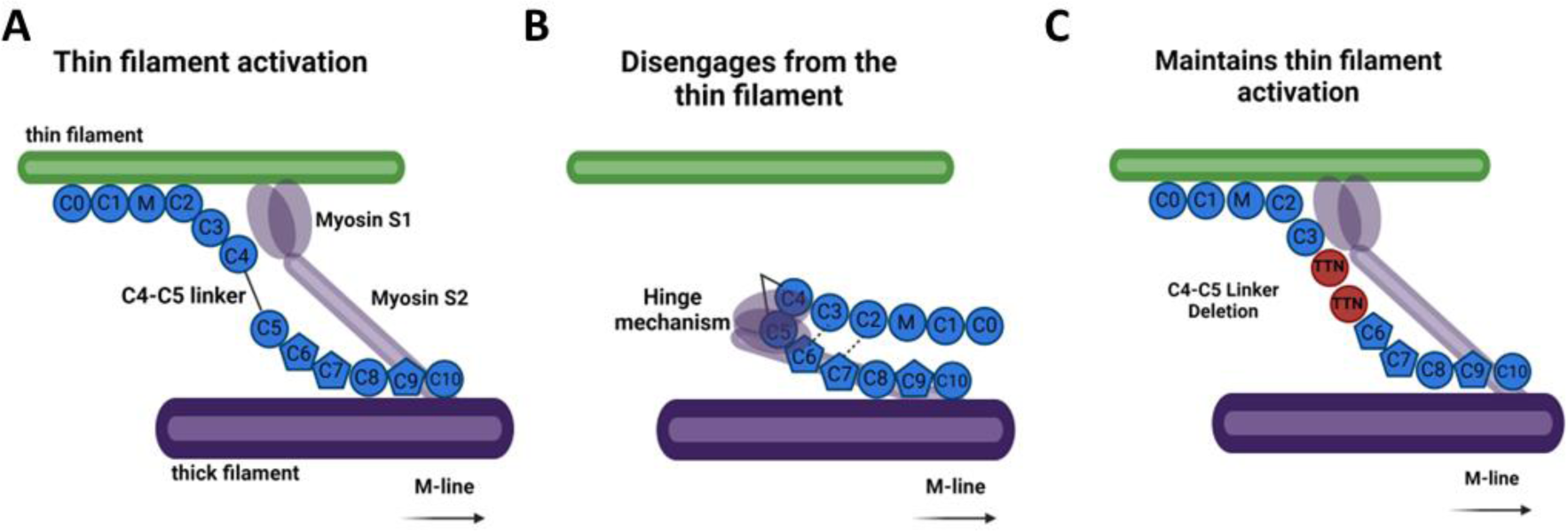
Proposed hinge model of cMyBP-C unbinding from the thin filament. Cartoon depiction of possible spatial arrangements of cMyBP-C with and without the middle domains. A) Cartoon showing cMyBP-C NTD bound to actin, aiding in thin filament activation and cross-bridge formation. B) NTD shown unbinding from actin due to the action of a hinge mechanism between domains C4 and C5, causing the NTD to move away spatially from the actin binding sites. Additionally, the hinge mechanism could explain how myosin S1 and S2 have high affinity for binding with the middle domains, which would help stabilize myosin into an “OFF” state and/or SRX state. Domains C3, C6 and/or C7 might help stabilize the bent form of the linker (denoted by dashed lines) and, thus, without these domains functionally present, there is still a high probability for the NTD to interact with actin and myosin heads to remain in an “ON” and/or disordered relaxed (DRX) state. C) Upon deletion of the C4-C5 linker from cMyBP-C (e.g., in C0C3-TTN_2_-C6C7-sc or ΔLinker-C0C7sc), the hinge cannot form and the NTD remain near or bound to actin, increasing the probability for activating the thin filament and increasing the probability of myosin heads in the “ON” and/or DRX states.

Alternatively, it is possible that the movement of the NTD in response to a conformational change in the C4-C5 linker might lead to alterations in interactions between myosin and cMyBP-C. Both the NTD^14,18^ and the middle domains^16,27–29^ can bind to myosin. Our data indicate that protein constructs that exclude the C4-C5 linker region cause a large increase in submaximal tension, which in principle could be explained by an increase in the proportion of myosin heads in the structural ON state. Previous work by others suggests that cMyBP-C is important in stabilizing the myosin mesa (portion of S1) onto the proximal portion of myosin S2^16^ suggesting that cMyBP-C helps stabilize myosin S1 into an OFF state and/or the lower-energy biochemical state, called the super-relaxed state (SRX)^47^. These data were recently confirmed using the fast skeletal paralog of MyBP-C, where cutting out endogenous MyBP-C^(C1-C7)^ with TEVp led to a significant shift towards the ON state in psoas muscle from SnoopC2 mice^38^. Without the middle domains or linker present, we found a hypercontractile state of muscle, which supports the hypothesis that the middle domains of cMyBP-C are responsible for stabilizing a certain proportion of myosin heads into the structural OFF state^38^ and/or the SRX state^29^.

We also showed that the middle domains are required for normal responses to stretch activation. Following a 2% stretch in a Ca^2+^ activated myocytes, the second phase (P2) of the stretch activation response was greatly affected in deletion and substitution protein constructs (i.e., C0C2sc, C0C2-TTN_5_-sc, C0C3-TTN_2_-C6C7-sc, C0C2-TTN-C4C5-TTN_2_-sc, and ΔLinker-C0C7sc). P2 is thought to represent the force decay that occurs after overstretching the cardiomyocyte and is likely a combination of elastic components and cross-bridge unbinding^37,48,49^. The third phase (P3) demonstrates the stretch activation phenomenon where there is a transient increase in tension beyond the pre-stretch steady state values^37^. In the linker deletion and spacer constructs, we found that there was a significant reduction in the total force decay during P2. Additionally, the new steady state force during P3 was augmented compared to before and after TEVp conditions. The overall shape of the stretch activation was significantly different due to the lack of force decay, i.e., increased P2 amplitude. Previous work in insect flight muscles suggested that the rise in force during P3 is due to rapid recruitment of non-force-generating cross-bridges into strongly bound, force-generating cross-bridges^50,51^. Computational modeling that simulated reduced cross-bridge activation resulted in a similar stretch activation response similar to that seen in our deletion and spacer constructs^48^. These results suggest that the middle domains play a role in limiting the amount of cross-bridges in strongly bound, force-generating states. Treatment with TEVp typically increased k*_rel_* consistent with previous findings^20^, but somewhat unexpected replacing cMyBP-C^C0-C7^ with spacer constructs and the ΔLinker-C0C7sc led to a slowing of force decay comparable to wild-type C0C7sc. This result may suggest that without the middle domains, or the C4-C5 linker, there is additional “drag” slowing cross-bridge detachment. Taken together, these studies suggest that the middle domains normally limit force-generative cross-bridges, potentially by limiting thin filament activation and/or the proportion of myosin heads in the ON state (Fig. 5B-C).

One limitation to this study is that the titin I27 domains inserted into the C0C7 cMyBP-C sequence could directly contribute to observed alterations in force mechanics. The I27 titin domain was chosen as a well-characterized yet unrelated Ig domain with similar size to the Ig domains found in cMyBP-C (∼4 nm) and with documented stability as a well-folded domain (i.e., requiring large atomic force microscopy forces to unfold the domains)^52^. However, others found that the I27 domains (also known as I91) can unfold and refold under more physiological forces^53^. To determine if the TTN domains directly affected the function of permeabilized cardiomyocytes, we tested the effects of replacing cMyBP-C^C0C7^ with TTN_5_-sc and found no effects on Ca^2+^ sensitivity, residual force, or stretch activation. These results support the idea that TTN spacer domains themselves did not significantly affect myocyte force or cross-bridge kinetics in our study.

In conclusion, we demonstrate the functional effects of the middle domains of cMyBP-C in Ca^2+^ activated, force generating sarcomeres. Specifically, deletion of these domains or substituting them with unrelated Ig domains led to significant increases in submaximal tension and abnormal responses to stretch. Importantly, the abnormal responses to stretch are consistent with prolonged crossbridge binding and slowed detachment suggesting that the middle domains of cMyBP-C are essential for normal relaxation. Results are thus consistent with an autoinhibitory role for these domains in the native full-length protein. We propose an autoinhibitory model where the middle domains fold via the C4-C5 linker hinge to interact with the N’-terminal domains thereby limiting the interactions of the N’-terminal domains (C0-C2) with its binding partners on thin or thick filaments.

## References

1. Heling, L. W. H. J., Geeves, M. A. & Kad, N. M. MyBP-C: one protein to govern them all. J. Muscle Res. Cell Motil. 41, 91–101 (2020).

2. Moss, R. L., Fitzsimons, D. P. & Ralphe, J. C. Cardiac MyBP-C regulates the rate and force of contraction in mammalian myocardium Cardiac Myosin Binding Protein C. Circ. Res. 116, 183–192 (2015).

3. Razumova, M. V. et al. Effects of the N-terminal domains of myosin binding protein-C in an in vitro motility assay: Evidence for long-lived cross-bridges. J. Biol. Chem. 281, 35846–35854 (2006).

4. Glazier, A. A., Thompson, A. & Day, S. M. Allelic imbalance and haploinsufficiency in MYBPC3-linked hypertrophic cardiomyopathy. Pflugers Arch. 471, 781–793 (2019).

5. Maron, B. J. Hypertrophic CardiomyopathyA Systematic Review. JAMA 287, 1308–1320 (2002).

6. Semsarian, C., Ingles, J., Maron, M. S. & Maron, B. J. New perspectives on the prevalence of hypertrophic cardiomyopathy. J. Am. Coll. Cardiol. 65, 1249–1254 (2015).

7. Harris, S. P., Lyons, R. G. & Bezold, K. L. In the Thick of It: HCM-Causing Mutations in Myosin Binding Proteins of the Thick Filament. Circ. Res. 108, 751–764 (2011).

8. Nadvi, N. A., Michie, K. A., Kwan, A. H., Guss, J. M. & Trewhella, J. Clinically Linked Mutations in the Central Domains of Cardiac Myosin-Binding Protein C with Distinct Phenotypes Show Differential Structural Effects. Struct. Lond. Engl. 1993 24, 105–115 (2016).

9. Razumova, M. V., Bezold, K. L., Tu, A.-Y., Regnier, M. & Harris, S. P. Contribution of the Myosin Binding Protein C Motif to Functional Effects in Permeabilized Rat Trabeculae. J. Gen. Physiol. 132, 575–585 (2008).

10. Shaffer, J. F., Kensler, R. W. & Harris, S. P. The myosin-binding protein C motif binds to F-actin in a phosphorylation-sensitive manner. J. Biol. Chem. 284, 12318–12327 (2009).

11. Kensler, R. W., Shaffer, J. F. & Harris, S. P. Binding of the N-terminal Fragment C0–C2 of Cardiac MyBP-C to Cardiac F-actin. J. Struct. Biol. 174, 44–51 (2011).

12. Whitten, A. E., Jeffries, C. M., Harris, S. P. & Trewhella, J. Cardiac myosin-binding protein C decorates F-actin: implications for cardiac function. Proc. Natl. Acad. Sci. U. S. A. 105, 18360–18365 (2008).

13. Kunst, G. et al. Myosin binding protein C, a phosphorylation-dependent force regulator in muscle that controls the attachment of myosin heads by its interaction with myosin S2. Circ. Res. 86, 51–58 (2000).

14. Gruen, M. & Gautel, M. Mutations in beta-myosin S2 that cause familial hypertrophic cardiomyopathy (FHC) abolish the interaction with the regulatory domain of myosin-binding protein-C. J. Mol. Biol. 286, 933–949 (1999).

15. Calaghan, S. C., Trinick, J., Knight, P. J. & White, E. A role for C-protein in the regulation of contraction and intracellular Ca2+ in intact rat ventricular myocytes. J. Physiol. 528, 151–156 (2000).

16. Nag, S. et al. The myosin mesa and the basis of hypercontractility caused by hypertrophic cardiomyopathy mutations. Nat. Struct. Mol. Biol. 24, 525–533 (2017).

17. Ababou, A., Gautel, M. & Pfuhl, M. Dissecting the N-terminal Myosin Binding Site of Human Cardiac Myosin-binding Protein C: STRUCTURE AND MYOSIN BINDING OF DOMAIN C2*. J. Biol. Chem. 282, 9204–9215 (2007).

18. Starr, R. & Offer, G. The interaction of C-protein with heavy meromyosin and subfragment-2. Biochem. J. 171, 813–816 (1978).

19. Inchingolo, A. V., Previs, S. B., Previs, M. J., Warshaw, D. M. & Kad, N. M. Revealing the mechanism of how cardiac myosin-binding protein C N-terminal fragments sensitize thin filaments for myosin binding. Proc. Natl. Acad. Sci. 116, 6828–6835 (2019).

20. Napierski, N. C. et al. A Novel ‘Cut and Paste’ Method for In Situ Replacement of cMyBP-C Reveals a New Role for cMyBP-C in the Regulation of Contractile Oscillations. Circ. Res. 126, 737–749 (2020).

21. Harris, S. P. et al. Hypertrophic Cardiomyopathy in Cardiac Myosin Binding Protein-C Knockout Mice. Circ. Res. 90, 594–601 (2002).

22. Wang, C., Schwan, J. & Campbell, S. G. Slowing of contractile kinetics by myosin-binding protein C can be explained by its cooperative binding to the thin filament. J. Mol. Cell. Cardiol. 96, 2–10 (2016).

23. de Lange, W. J., Grimes, A. C., Hegge, L. F. & Ralphe, J. C. Ablation of cardiac myosin–binding protein-C accelerates contractile kinetics in engineered cardiac tissue. J. Gen. Physiol. 141, 73–84 (2013).

24. Flashman, E., Watkins, H. & Redwood, C. Localization of the binding site of the C-terminal domain of cardiac myosin-binding protein-C on the myosin rod. Biochem. J. 401, 97–102 (2007).

25. Gilbert, R., Cohen, J. A., Pardo, S., Basu, A. & Fischman, D. A. Identification of the A-band localization domain of myosin binding proteins C and H (MyBP-C, MyBP-H) in skeletal muscle. J. Cell Sci. 112 **(Pt** **1**), 69–79 (1999).

26. Gilbert, R., Kelly, M. G., Mikawa, T. & Fischman, D. A. The carboxyl terminus of myosin binding protein C (MyBP-C, C-protein) specifies incorporation into the A-band of striated muscle. J. Cell Sci. 109 **(Pt** **1**), 101–111 (1996).

27. Dutta, D., Nguyen, V., Campbell, K. S., Padrón, R. & Craig, R. Cryo-EM structure of the human cardiac myosin filament. Nature 623, 853–862 (2023).

28. Tamborrini, D. et al. Structure of the native myosin filament in the relaxed cardiac sarcomere. Nature 623, 863–871 (2023).

29. Ponnam, S. & Kampourakis, T. Microscale thermophoresis suggests a new model of regulation of cardiac myosin function via interaction with cardiac myosin-binding protein C. J. Biol. Chem. 298, 101485 (2022).

30. Doh, C. Y. et al. Molecular characterization of linker and loop-mediated structural modulation and hinge motion in the C4-C5 domains of cMyBPC. J. Struct. Biol. 214, 107856 (2022).

31. Idowu, S. M., Gautel, M., Perkins, S. J. & Pfuhl, M. Structure, stability and dynamics of the central domain of cardiac myosin binding protein C (MyBP-C): implications for multidomain assembly and causes for cardiomyopathy. J. Mol. Biol. 329, 745–761 (2003).

32. Guardiani, C., Cecconi, F. & Livi, R. Computational analysis of folding and mutation properties of C5 domain of myosin binding protein C. Proteins 70, 1313–1322 (2008).

33. Rybakova, I. N., Greaser, M. L. & Moss, R. L. Myosin binding protein C interaction with actin: characterization and mapping of the binding site. J. Biol. Chem. 286, 2008–2016 (2011).

34. Colson, B. A., Rybakova, I. N., Prochniewicz, E., Moss, R. L. & Thomas, D. D. Cardiac myosin binding protein-C restricts intrafilament torsional dynamics of actin in a phosphorylation-dependent manner. Proc. Natl. Acad. Sci. U. S. A. 109, 20437–20442 (2012).

35. Zakeri, B., et al. Peptide tag forming a rapid covalent bond to a protein, through engineering a bacterial adhesin. Proc. Natl. Acad. Sci. 109, E690–E697 (2012).

36. van Dijk, S. J. et al. Point mutations in the tri-helix bundle of the M-domain of cardiac myosin binding protein-C influence systolic duration and delay cardiac relaxation. J. Mol. Cell. Cardiol. 119, 116–124 (2018).

37. Stelzer, J. E., Dunning, S. B. & Moss, R. L. Ablation of cardiac myosin-binding protein-C accelerates stretch activation in murine skinned myocardium. Circ. Res. 98, 1212–1218 (2006).

38. Hessel, A. L. et al. Myosin-binding protein C regulates the sarcomere lattice and stabilizes the OFF states of myosin heads. Nat. Commun. 15, 2628 (2024).

39. Lee, K., Harris, S. P., Sadayappan, S. & Craig, R. Orientation of myosin binding protein C in the cardiac muscle sarcomere determined by domain-specific immuno-EM. J. Mol. Biol. 427, 274–286 (2015).

40. Risi, C. M. et al. Cryo-Electron Microscopy Reveals Cardiac Myosin Binding Protein-C M-Domain Interactions with the Thin Filament. J. Mol. Biol. 434, 167879 (2022).

41. Ratti, J., Rostkova, E., Gautel, M. & Pfuhl, M. Structure and Interactions of Myosin-binding Protein C Domain C0. J. Biol. Chem. 286, 12650–12658 (2011).

42. Moolman-Smook, J. et al. Identification of novel interactions between domains of Myosin binding protein-C that are modulated by hypertrophic cardiomyopathy missense mutations. Circ. Res. 91, 704–711 (2002).

43. Dominic, K. L. et al. The contribution of N-terminal truncated cMyBPC to in vivo cardiac function. J. Gen. Physiol. 155, e202213318 (2023).

44. Previs, M. J. et al. Phosphorylation and calcium antagonistically tune myosin-binding protein C’s structure and function. Proc. Natl. Acad. Sci. 113, 3239–3244 (2016).

45. Luther, P. K. et al. Direct visualization of myosin-binding protein C bridging myosin and actin filaments in intact muscle. Proc. Natl. Acad. Sci. U. S. A. 108, 11423–11428 (2011).

46. Flavigny, J. et al. Biomolecular interactions between human recombinant beta-MyHC and cMyBP-Cs implicated in familial hypertrophic cardiomyopathy. Cardiovasc Res 60, 388–96 (2003).

47. McNamara, J. W. et al. Ablation of cardiac myosin binding protein-C disrupts the super-relaxed state of myosin in murine cardiomyocytes. J. Mol. Cell. Cardiol. 94, 65–71 (2016).

48. Campbell, K. B. & Chandra, M. Functions of Stretch Activation in Heart Muscle. J. Gen. Physiol. 127, 89–94 (2006).

49. Jarvis, K. J., Bell, K. M., Loya, A. K., Swank, D. M. & Walcott, S. Force-velocity and tension transient measurements from Drosophila jump muscle reveal the necessity of both weakly-bound cross-bridges and series elasticity in models of muscle contraction. Arch. Biochem. Biophys. 701, 108809 (2021).

50. Getz, E. B., Cooke, R. & Lehman, S. L. Phase transition in force during ramp stretches of skeletal muscle. Biophys. J. 75, 2971–2983 (1998).

51. Linari, M., Reedy, M. K., Reedy, M. C., Lombardi, V. & Piazzesi, G. Ca-activation and stretch-activation in insect flight muscle. Biophys. J. 87, 1101–1111 (2004).

52. Li, H. et al. Reverse engineering of the giant muscle protein titin. Nature 418, 998–1002 (2002).

53. Rivas-Pardo, J. A. et al. Work done by titin protein folding assists muscle contraction. Cell Rep. 14, 1339–1347 (2016).

