## Supplemental Materials for "Autoinhibition of cMyBP-C by its middle domains"

### Methods

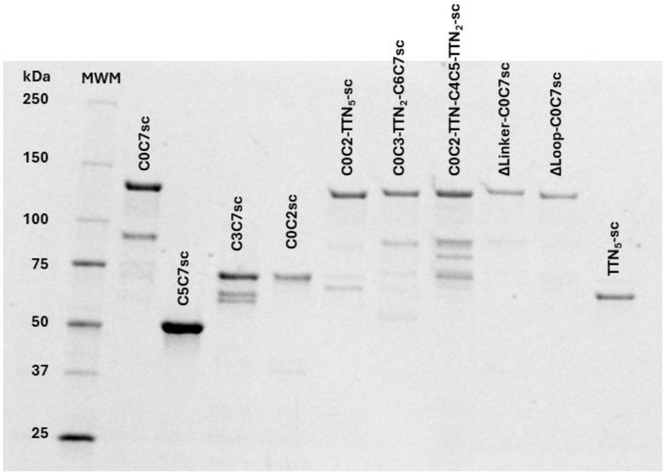


**Supplement Fig. S1:** Stain-free gel of final proteins used for force mechanics experiments.

### Results

#### Replacement of cMyBP-C$^{C0C7}$ with TTN$_5$-sc

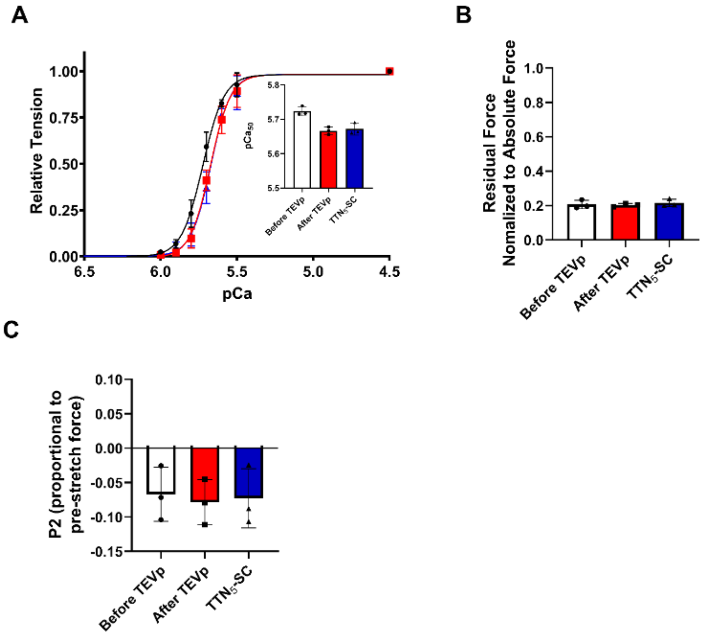

**Supplement Fig. S2:** Force mechanics data of the $TTN_5$-sc construct. A) Tension-pCa curves before TEVp (black), after TEVp treatment (red), and after pasting $TTN_5$-sc (blue). B) Residual force analysis between the three conditions. C) P2 amplitude relative to the pre-stretch force across the three conditions.

### Hill Slope Analysis

| ***Conditions*** | | **Hill Slope** |
|---|---|---|
| **C0C7sc** | **Before TEVp** | 3.76 ± 0.52 |
| | **After TEVp** | 3.63 ± 0.46 |
| | **C0C7sc** | 4.47 ± 0.71 * |
| **C5C7sc** | **Before TEVp** | 5.00 ± 0.85 |
| | **After TEVp** | 4.43 ± 1.11 |
| | **C5C7sc** | 4.31 ± 0.66 |
| **C3C7sc** | **Before TEVp** | 6.47 ± 0.44 |
| | **After TEVp** | 4.38 ± 0.47 ‡‡ |
| | **C3C7sc** | 4.74 ± 0.94 * |
| **C0C2sc** | **Before TEVp** | 4.34 ± 0.50 |
| | **After TEVp** | 4.29 ± 0.42 |
| | **C0C2sc** | 5.21 ± 1.38 |
| **C0C2-$TTN_5$-sc** | **Before TEVp** | 9.50 ± 1.87 |
| | **After TEVp** | 8.13 ± 3.22 |
| | **C0C2-$TTN_5$-sc** | 2.00 ± 0.36 ** |
| **C0C3-$TTN_2$-C6C7-sc** | **Before TEVp** | 5.01 ± 0.95 |
| | **After TEVp** | 4.71 ± 0.69 |
| | **C0C3-$TTN_2$-C6C7-sc** | 2.98 ± 0.43 * |
| **C0C2-TTN-C4C5-$TTN_2$-sc** | **Before TEVp** | 7.38 ± 1.41 |
| | **After TEVp** | 6.92 ± 2.09 |
| | **C0C2-TTN-C4C5-$TTN_2$-sc** | 2.37 ± 0.40 *** |
| **ΔLinker-C0C7sc** | **Before TEVp** | 6.70 ± 1.63 |
| | **After TEVp** | 6.22 ± 1.28 |
| | **ΔLinker-C0C7sc** | 3.36 ± 0.57 ** |
| **ΔLoop-C0C7sc** | **Before TEVp** | 6.97 ± 1.28 |
| | **After TEVp** | 6.40 ± 1.35 |
| | **ΔLoop-C0C7sc** | 7.19 ± 1.59 |

**Supplemental Table SI Legend**: Average ± SD of hill slopes before, after TEVp, and after protein replacement. * = $p < 0.05$ Before TEVp vs. after protein replacement. ** = $p < 0.01$ Before TEVp vs. after protein replacement. *** = $p < 0.001$ Before TEVp vs. after protein replacement. ‡‡ = $p < 0.01$ Before TEVp vs After TEVp.

### $k_{df}$ Analysis

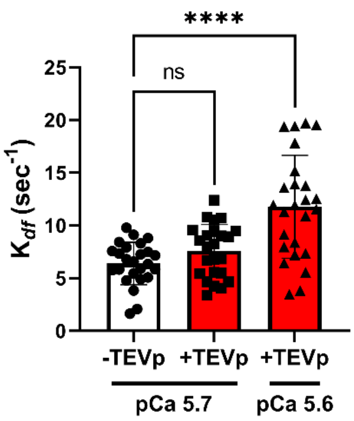


**Supplemental Fig. S3:** The rate of delayed force redevelopment after a rapid stretch, or $k_{df}$, was compared before (-TEVp) and after TEVp (+TEVp) conditions at submaximal pCa 5.7 or 5.6. The rates fit from C0C2sc, C0C2-$TTN_5$-sc, C0C3-$TTN_2$-C6C7-sc, C0C2-TTN-C4C5-$TTN_2$-sc, and ΔLinker-C0C7sc were collated for -TEVp and +TEVp due to the inability to fit a single exponential curve to the post-protein conditions in all five of the above protein constructs. Due to lower tension with +TEVp (i.e., rightward shift in the tension-pCa relationship), we compared -TEVp to +TEVp at pCa 5.7 and 5.6. A one-way ANOVA was used to determine significance, denoted as **** < 0.0001.

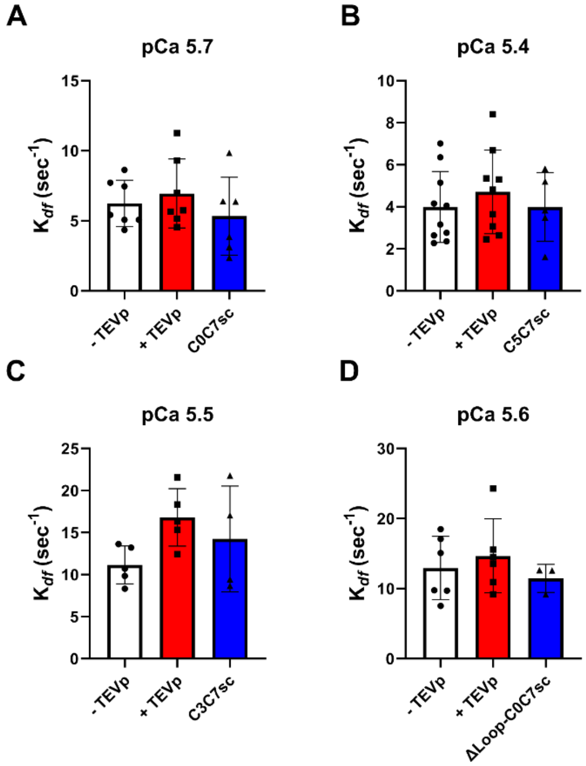


**Supplemental Fig. S4:** $k_{df}$ at submaximal calcium concentrations were compared before TEVp treatment (-TEVp), after TEVp treatment (+TEVp), and after pasting A) C0C7sc, B) C5C7sc, C) C3C7sc, and D) ΔLoop-C0C7sc. A repeated measures one-way ANOVA was used to compare the three conditions and there no significant differences across groups.
